# Molecular mechanisms of Cardiac Injury associated with myocardial SARS-CoV-2 infection

**DOI:** 10.1101/2020.07.27.220954

**Authors:** XianFang Liu, LongQuan Lou, Lei Zhou

**Author notes:** those authors contributed equally to this work (XianFang Liu, LongQuan Lou). Correspondence: Lei Zhou, Department of Cardiology, The First Affiliated Hospital of Nanjing Medical University, 300 Guangzhou Road, Nanjing, 210029, Jiangsu, China.

## Abstract

**Background:** Coronavirus Disease 2019 (COVID-19) caused by severe acute respiratory syndrome coronavirus 2 (SARS-CoV-2) has spread around the world. Developing cardiac injury is a common condition in COVID-19 patients, but the pathogenesis remains unclear.

**Methods:** The RNA-Seq dataset (GES150392) compared expression profiling of mock human induced pluripotent stem cell-derived cardiomyocytes (hiPSC-CMs) and SARS-cov-2-infected hiPSC-CMs were obtained from Gene Expression Omnibus (GEO). We identified the differentially expressed genes (DEGs) between those two groups. Through gene set enrichment analysis (GSEA), Gene Ontology (GO) analysis, Kyoto Encyclopedia of Genes and Genomes (KEGG) pathway analysis, and CLINVAR human diseases analysis to identify the main effect of SARS-CoV-2 on cardiomyocytes. A protein-protein interaction (PPI) network was constructed to visualize interactions and functions of the hub genes.

**Results:** A total of 1554 DEGs were identified (726 upregulated genes and 828 downregulated genes). Gene enrichment analysis shown that SARS-CoV-2 activate immuno-inflammatory responses via multiple signal pathways, including TNFα, IL6-JAK-STAT3, IL2-STAT5, NF-κB, IL17, and Toll-like receptor signaling pathway in hiPSC-CMs. Whereas, the muscle contraction, cellular respiration and cell cycle of hiPSC-CMs were inhibited by SARS-CoV-2. CLINVAR human diseases analysis shown SRAS-Cov-2 infection was associated with myocardial infarction, cardiomyopathy and Limb-girdle muscular dystrophy. 15 hub genes were identified based on PPI network. Function analysis revealed that 11 upregulated hub genes were mainly enriched in cytokine activity, chemokine activity, Inflammatory response, leukocyte chemotaxis, and lipopolysaccharide-mediated signaling pathway. Furthermore, 4 downregulated hub genes were related to cell cycle regulation.

**Conclusion:** The present study elucidates that the SARS-CoV-2 infection induced a strong defensive response in cardiomyocyte, leading to excessive immune inflammation, cell hypoxia, functional contractility reduction and apoptosis, ultimately result in myocardial injury.

## Introduction

Coronavirus disease 2019 (COVID-19) is a viral pandemic caused by the severe acute respiratory syndrome coronavirus 2 (SARS-CoV-2)[1]. In December 2019, COVID-19 was first reported in Wuhan of China as a severe unknown form of pneumonia. Subsequently, a global pandemic was declared by World Health Organization (WHO) in March 2020. At the time of preparing this manuscript, there was more than 500,000 fatalities caused by COVID-19.

Acute myocardial damage is the most common described cardiovascular (CV) complication in COVID-19 patients [2], In multivariable adjusted models, cardiac injury has been identified as a significantly and independently risk factor (hazard ratios=4.26) associated with mortality[3]. Cytokine storms (caused by acute systemic inflammation)[1, 4], hypoxemia[5], and pathogen-mediated damage [6, 7]were considered as the potential mechanisms responsible for CV complications in COVID-19. However, the exact mechanism of SARS-CoV-2 infection-related myocardial injury remains unclear.

Arun Sharma et al reported that SARS-cov-2 directly infected human induced pluripotent stem cell-derived cardiomyocytes (hiPSC-CMs) in vitro, induce contractility depletion and apoptosis[8]. They obtain the expression profiling of 3 SARS-CoV-2 Infected hiPSC-CMs samples and 3 Mock hiPSC-CMs samples by high throughput sequencing, and deposit it on the National Center for Biotechnology Information (NCBI) Gene Expression Omnibus (GEO) database (GSE150392). In this study, we performed a detailed bioinformatics analysis of the GSE150392 RNA-seq data to further examine the specific mechanisms of myocardial damage caused by SARS-CoV-2.

## Materials and methods

### 1. Data sources

We obtained the RNA-seq data (GSE150392) from the National Center for Biotechnology Information (NCBI) Gene Expression Omnibus (GEO) database. The GSE150392 dataset has a total of 6 samples, containing 3 SARS-CoV-2 Infected hiPSC-CMs (human induced pluripotent stem cell-derived cardiomyocytes) samples (cov1-3: GSM4548303-5) and Mock hiPSC-CMs samples (mock1-3: GSM4548306-8).

### 2. Identification of differentially expressed genes (DEGs)

We used R (version 4.0.1, https://www.R-project.org/) package DESeq2 (version 1.28.1) to determine DEGs [9] (P adj < 0.01, | (log2FoldChange| > 2) between SARS-CoV-2 Infected hiPSC-CMs and Mock hiPSC-CMs. Heatmap of DEGs was generated with R pheatmap package (version 1.0.12, https://CRAN.R-project.org/package=pheatmap).

### 3. Gene Set Enrichment Analysis (GSEA) of all detected genes

GSEA was performed using the GSEA software (version 4.0.3) implementation in our study for identifying potential hallmark of SARS-CoV-2 Infected hiPSC-CMs[10]. The annotated gene sets of h.all.v7.1.symbols.gmt were adopted from The Molecular Signatures Database (MSigDB, http://www.broad.mit.edu/gsea/msigdb/index.jsp). We performed 1,000 times of permutations. Collapse dataset to gene symbols was “False.” The permutation type was “gene set.”

### 4. Enrichment analyses of DEGs

Gene Ontology (GO) analysis (biological processes, molecular function and cellular component) and Kyoto Encyclopedia of Genes and Genomes (KEGG) pathway enrichment were analyzed using R package clusterProfiler (version 3.16.0)[11]. GO analysis (Immune System Process), KEGG pathways functionally grouped networks and CLINVAR human diseases analysis using the CluGO (version 2.5.7) [12]and CluePedia (version 1.5.7) [13]apps of Cytoscape Software (Version3.8.0)[14].

### 5. Protein-Protein Interaction (PPI) Network

The online tool of Search tool for the retrieval of interacting genes (STRING, https://string-db.org) [15]was applied to establish a PPIs of DEGs with score (median confidence) >0.4. Cytoscape and CytoHubba (Version 0.1) [16]was used to visualized the PPI network and identify hub genes. GeneMANIA online database (https://genemania.org/) was used to analysis the hub genes[17].

## Results

### 1. SRAS-CoV-2 infection caused a large number of gene expression changes in hiPSC-CMs

The boxplot of expression values of all transcripts indicated similar whole transcriptome expression in each sample (Fig. 1A) and sample to sample distance heatmap revealed a relatively clear distinction between samples in the two groups (Fig. 1B). Based on the criteria of P < 0.01, and |logFC|>2(Fig.1C), a total of 1554 DEGs was identified from GSE150392, including 726 upregulated genes and 828 downregulated genes (Fig.1D).

**Fig 1:**
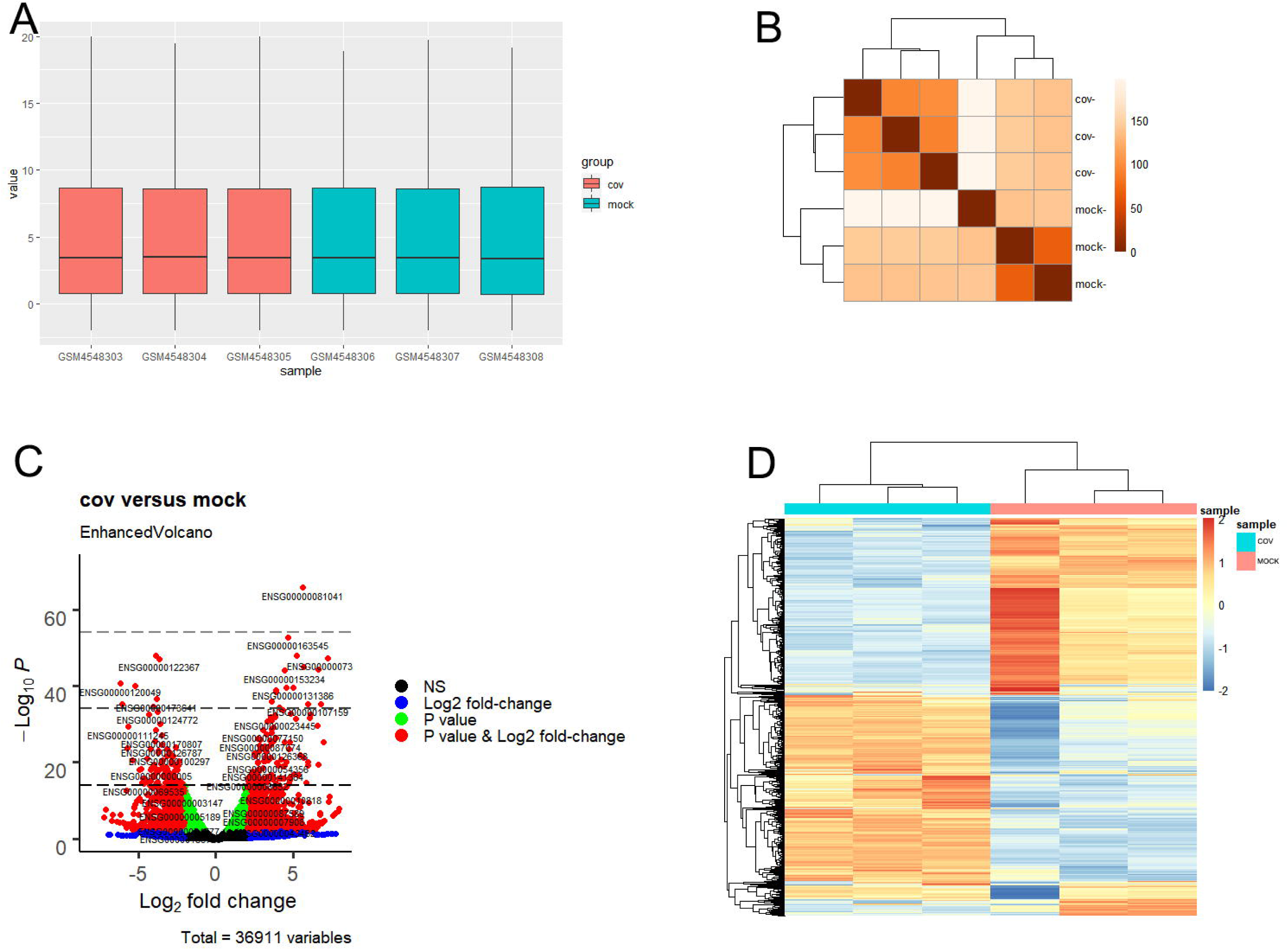
Differential expression of data between two sets of samples. (A) Standardization of gene expression; (B) sample to sample distance heatmap of all samples in COV and MOCK group; (C) Volcano plot of genes (The red points represent genes screened on the basis of |fold change| >2.0 and a corrected P-value of <0.01.); (D) Heat map of 1554 representative DEGs (Red areas represent highly expressed genes and green areas represent lowly expressed genes. Their phylogenetic relationships were shown on the left tree. The top tree indicated the cluster relationship of the samples). NS: no significant difference.

### 2. GSEA shown SRAS-CoV-2 infection was mainly associated with immune inflammatory response activity

To obtain insight into the effect of SARS-cov-2 to the heart, GSEA was used to map into hallmarks, 14 significant gene sets were shown (Table1, Fig.2A-N). GSEA demonstrated that many inflammation-related gene sets, such as TNFα signaling via NF-κB, interferon-γ response, interferon-α response, inflammatory response, IL6-JAK-STAT3 signaling, IL2-STAT5 signaling, hypoxia, P53 pathway and apoptosis gene sets were positively enriched by SARS-CoV-2 infection in hiPSC-CMs. Oxidative phosphorylation, E2F targets, G2M checkpoint, myogenesis and MYC targets v1 gene sets were negatively enriched by SARS-CoV-2 infection in hiPSC-CMs, which were associated with cell cycle, aerobic respiration and myogenesis.

**Table 1.** The most significant gene sets of COV vs Mock in GSEA.

| Gene Set follow link to MSigDB | ES | NES | FDR q-val |
| --- | --- | --- | --- |
| HALLMARK_TNFA_SIGNALING_VIA_NFKB | 0.694622 | 2.809924 | 0 |
| HALLMARK_INTERFERON_GAMMA_RESPONSE | 0.666493 | 2.694757 | 0 |
| HALLMARK_INTERFERON_ALPHA_RESPONSE | 0.702799 | 2.559929 | 0 |
| HALLMARK_INFLAMMATORY_RESPONSE | 0.623106 | 2.533835 | 0 |
| HALLMARK_IL6_JAK_STAT3_SIGNALING | 0.592646 | 2.153886 | 0 |
| HALLMARK_IL2_STAT5_SIGNALING | 0.454536 | 1.847625 | 4.36E-04 |
| HALLMARK_HYPOXIA | 0.429023 | 1.733556 | 0.001023 |
| HALLMARK_P53_PATHWAY | 0.429149 | 1.731462 | 9.44E-04 |
| HALLMARK_APOPTOSIS | 0.399164 | 1.570103 | 0.0065 |
| HALLMARK_OXIDATIVE_PHOSPHORYLATION | -0.59549 | -2.48047 | 0 |
| HALLMARK_E2F_TARGETS | -0.55286 | -2.29242 | 0 |
| HALLMARK_G2M_CHECKPOINT | -0.47966 | -1.99993 | 0 |
| HALLMARK_MYOGENESIS | -0.41285 | -1.70035 | 0 |
| HALLMARK_MYC_TARGETS_V1 | -0.37378 | -1.56712 | 0.002331 |

(ES: enrichment score; NES: normalized enrichment score FDR: false discovery rate.)

### 3. GO term enrichment analyses and CLINVAR human diseases analysis of DEGs

GO analysis of DEGs was divided into four functional groups, including biological processes (BP), cell composition (CC), molecular function (MF) and Immune System Process (ISP). The top 5 results of BP, CC and MF are shown in Fig.3.

In the BP group, the upregulated genes were mainly enriched in response to virus, response to lipopolysaccharide, response to molecule of bacterial origin, leukocyte cell-cell adhesion and defense response to virus. The downregulated genes were mainly concentrated in muscle system process, muscle contraction, oxidative phosphorylation, striated muscle contraction and respiratory electron transport chain.

In the CC group the upregulated genes were mainly enriched in receptor complex. The downregulated genes were mainly concentrated in myofibril, contractile fiber, sarcomere, I band and inner mitochondrial membrane protein complex.

In the MF group, the upregulated genes were mainly involved in terms about cytokine receptor binding, cytokine activity, DNA-binding transcription activator activity RNA polymerase II-specific, DNA-binding transcription activator activity and chemokine activity. The downregulated genes were mainly enriched in actin binding, NADH dehydrogenase activity, NADH dehydrogenase (ubiquinone) activity, NADH dehydrogenase (quinone) activity and structural constituent of muscle.

The ISP results were shown as Table 2. Type I interferon signaling pathway, regulation of adaptive immune response, neutrophil chemotaxis, regulation of type 2 immune response and CD4-positive α-β T cell cytokine production were enriched by upregulated genes. Positive regulation of megakaryocyte differentiation, regulation of megakaryocyte differentiation, megakaryocyte differentiation and thymus development were enriched by downregulated genes. CLINVAR human diseases analysis indicate that DEGs are significantly involved in terms about myocardial infarction-1, cardiomyopathy and Limb-girdle muscular dystrophy. (Table.3)

**Table 2.** GO (Immune System Process) analysis of DEGs.

| Term | Description | P Value |
| --- | --- | --- |
| Upregulated Genes |  |  |
| GO:0060337 | type I interferon signaling pathway | 4.27E-11 |
| GO:0002819 | regulation of adaptive immune response | 1.37E-06 |
| GO:0030593 | neutrophil chemotaxis | 1.05E-05 |
| GO:0002828 | regulation of type 2 immune response | 1.35E-05 |
| GO:0035743 | CD4-positive, alpha-beta T cell cytokine production | 6.05E-05 |
| Downregulated Genes |  |  |
| GO:0045654 | positive regulation of megakaryocyte differentiation | 2.89E-04 |
| GO:0045652 | regulation of megakaryocyte differentiation | 7.53E-04 |
| GO:0030219 | megakaryocyte differentiation | 0.002297 |
| GO:0048538 | thymus development | 0.033605 |

**Table 3.** CLINVAR human diseases analysis of DEGs.

| Term | Description | P Value |
| --- | --- | --- |
| Upregulated Genes |  |  |
| C1832662 | Myocardial infarction1 | 0.007240 |
| Downregulated Genes |  |  |
| C0878544 | Cardiomyopathy | 1.01E-15 |
| C0686353 | Limb-girdle muscular dystrophy | 9.51E-05 |

### 4. KEGG enrichment analysis of DEGs

The signaling pathways of DEGs were shown in Fig.4. The data was imported into Cytoscape to calculate the topological characteristics of the network and determine each node. Cytokine-cytokine receptor interaction, TNF signaling pathway, influenza A, NF-κB signaling pathway, osteoclast differentiation, measles, viral protein interaction with cytokine & cytokine receptor, IL-17 signaling pathway, Toll-like receptor signaling pathway and rheumatoid arthritis were significantly enriched by upregulated DEGs (Fig.4A). Thermogenesis, parkinson disease, oxidative phosphorylation, huntington disease, cardiac muscle contraction, non-alcoholic fatty liver disease (NAFLD), retrograde endocannabinoid signaling, adrenergic signaling in cardiomyocytes, dilated cardiomyopathy (DCM) and hypertrophic cardiomyopathy (HCM) were significantly enriched by downregulated DEGs (Fig.4B). The top 7 p-value KEGG pathways with its target genes were shown (Fig.4C).

### 5. PPI network analysis and hub genes recognition

The top 15 genes with the highest interaction degrees were identified, including 11 upregulated genes and 4 downregulated genes (Table 4). Then those genes were used for constructing the PPI network (Fig.2A). 11 upregulated hub genes showed the complex PPI network with the Co-expression of 66.42%, Co-localization of 9.39%, Genetic interactions of 8.72%, Predicted of 8.60%, Shared protein domains of 3.99%, pathway of 1.8% and Physical Interactions of 1.09% (Fig. 2C). cytokine activity, chemokine activity, Inflammatory response, leukocyte chemotaxis and lipopolysaccharide-mediated signaling pathway were identified as the main function of those genes. 4 downregulated hub genes showed the complex PPI network with the Physical Interactions of 67.64%, Co-expression of 13.50%, Predicted of 6.35%, Co-localization of 6.17%, pathway of 4.35%, Genetic interactions of 1.4%, and Shared protein domains of 0.59% (Fig. 2b). Those genes were related to mitosis, nuclear division, metaphase/anaphase transition of cell cycle and regulation of ubiquitin-protein ligase activity.

**Table 4.**
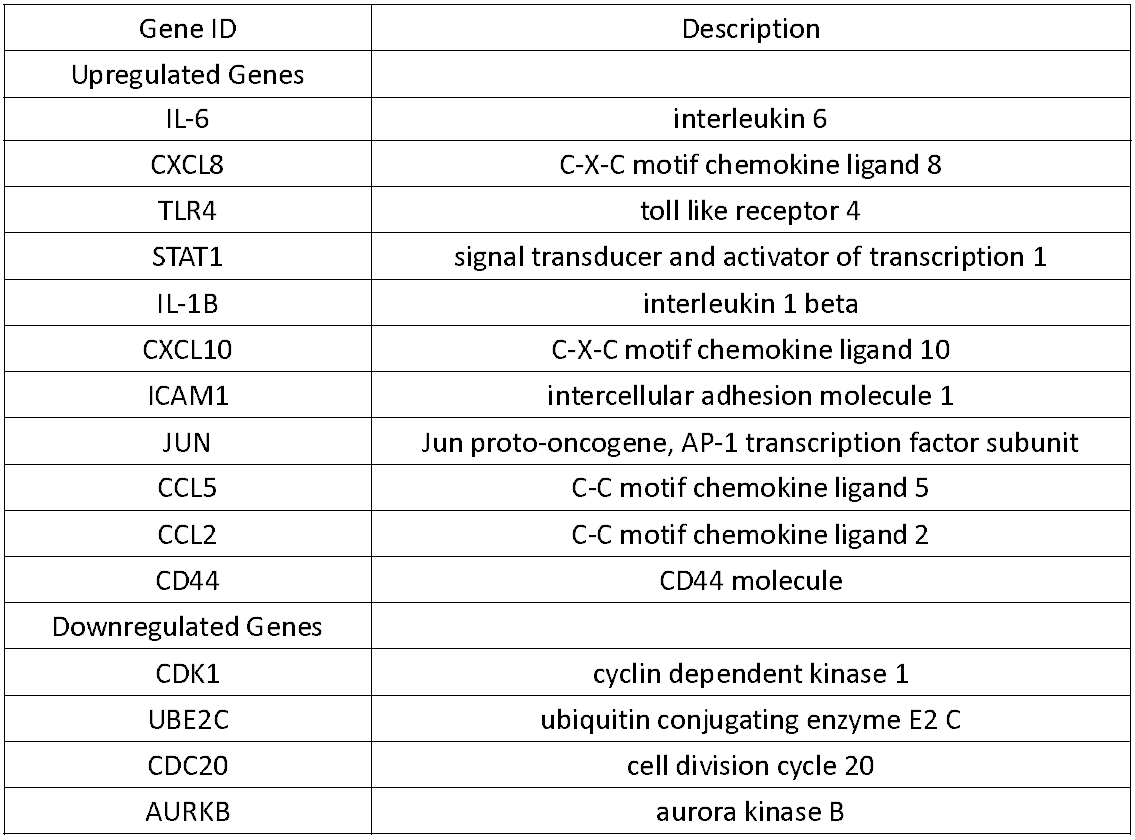
Top 15 hub genes with the highest interaction degrees in PPI network analysis.

**Fig 2:**
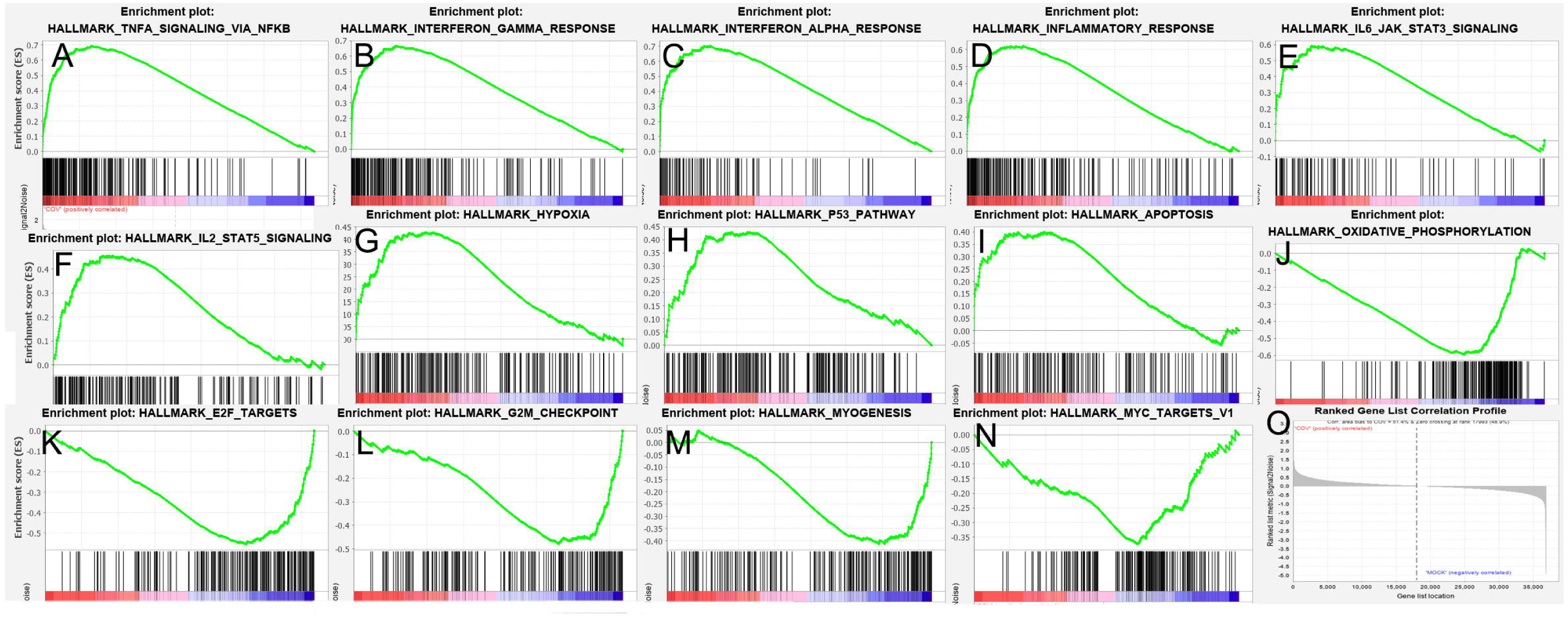
Gene set enrichment analysis (GSEA) of two groups (cov vs mock). 14 representative gene sets were listed. (A-I): Positive gene sets enriched in Cov vs Mock GSEA analysis; (J-N) Negative gene sets enriched in Cov vs Mock GSEA analysis; (O) Ranked Gene List Correlation Profile.

**Fig 3:**
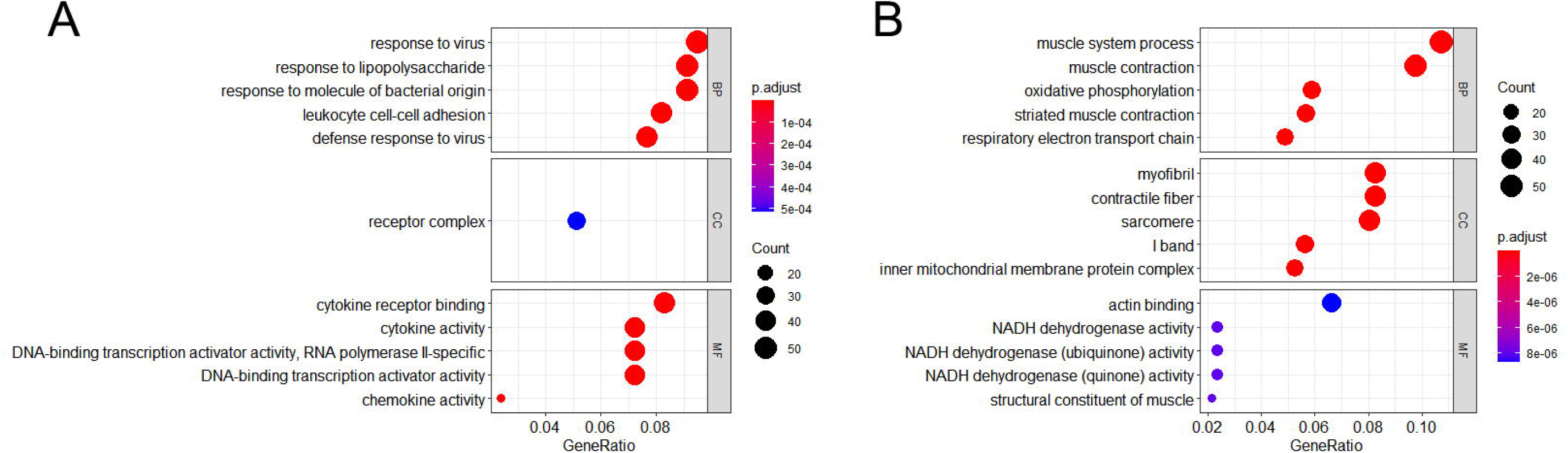
GO analysis (BP, CC and MF) of DEGs. (A) GO terms of upregulated DEGs. (B) GO terms of downregulated DEGs.

**Fig 4:**
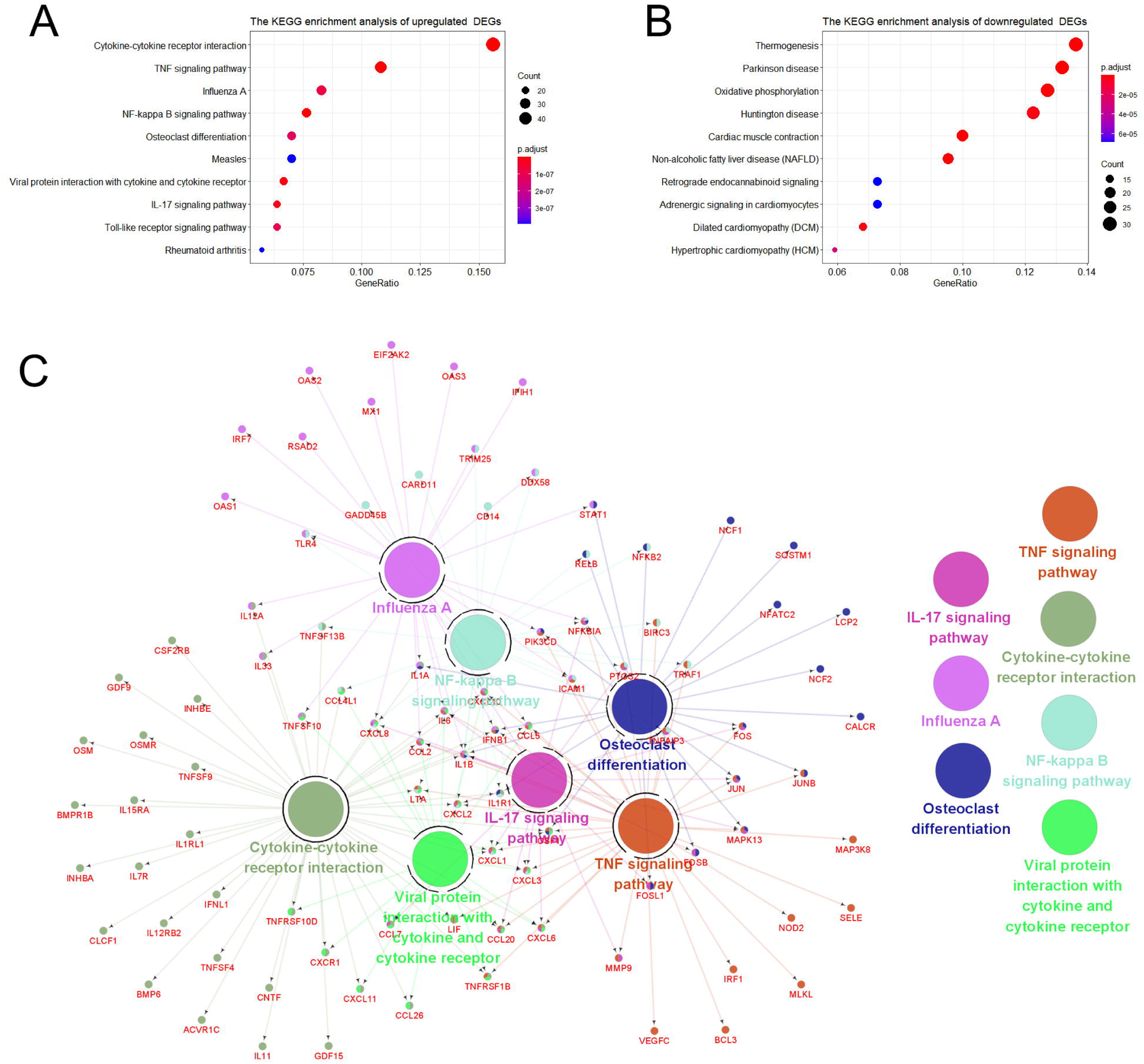
the KEGG enrichment analysis of DEGs. (A) KEGG pathways of upregulated DEGs. (B) KEGG pathways of downregulated DEGs. (C) Representative pathways to genes network of upregulated DEGs, validated genes (in red) targeted by KEGG pathways.

**Fig 5:**
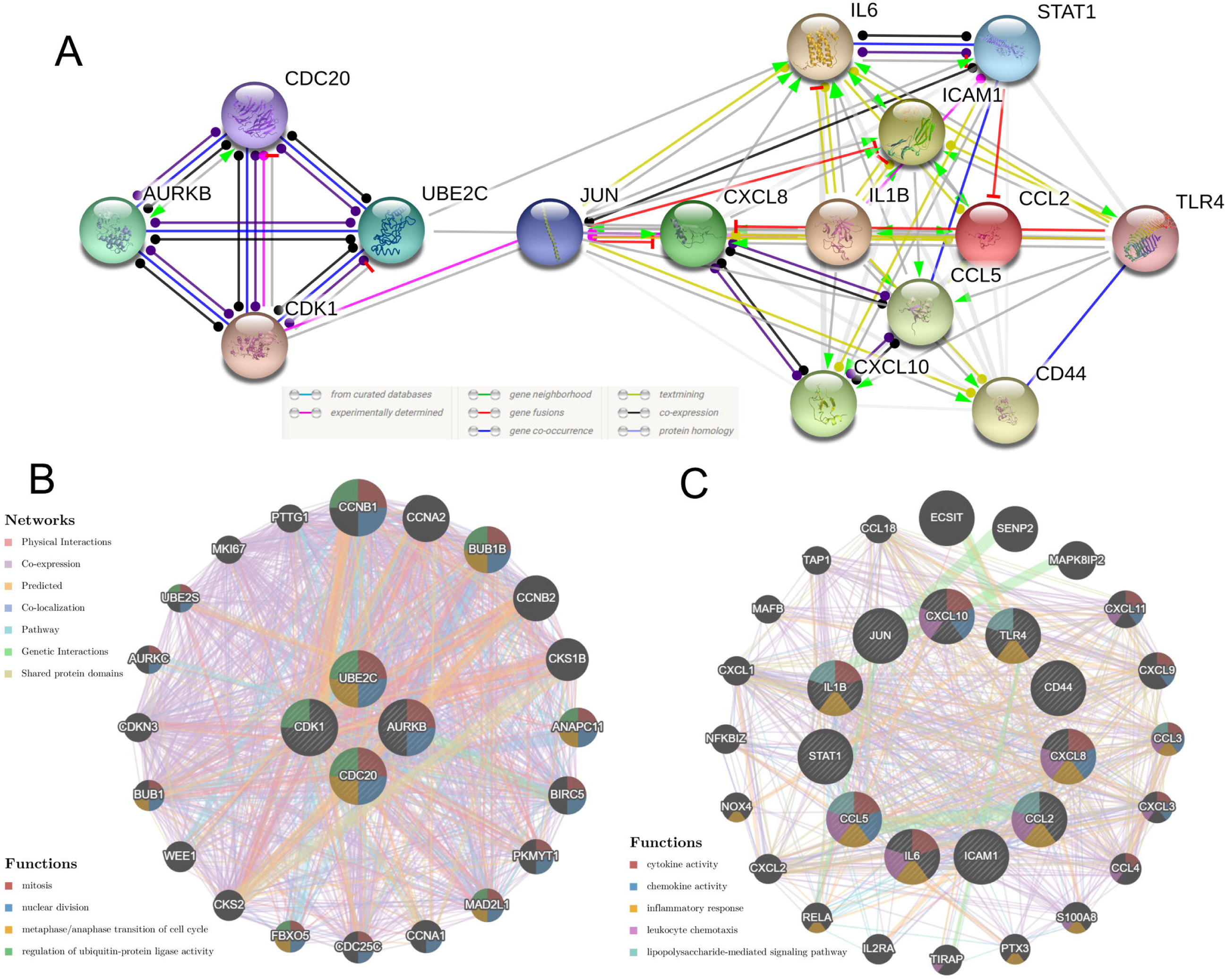
PPI network of Hub Genes. (A) The PPI network of the top 15 hub genes created by STRING; (B) PP1 networks and function analyses of the 4 downregulated hub genes; (C) PPI networks and function analyses of the 11 upregulated hub genes

## Discussion

SARS-CoV-2, SARS-CoV and MERS-CoV (Middle East respiratory syndrome coronavirus) are enveloped positive RNA viruses, belong to the Coronaviridae family β genera [18, 19]. SARS-CoV-2 and SARS-CoV utilize the same receptor angiotensin-converting enzyme 2 (ACE2) for invading human bodies. Those coronaviruses spread widely among people and lead to a potentially fatal disease [20, 21]. COVID-19 caused by SARS-CoV-2 has been spreading in 216 countries/ areas/ territories, with over 12,000,000 confirmed cases (World Health Organization statistics as on July 13, 2020) [22].

The Cardiac injury in COVID-19 was generally defined as the elevation of cardiac-specific biomarkers (eg, troponin concentration above the 99^th^ percentile upper reference limit) [1, 2]. It occurs in roughly 8-13% [23, 24]in confirmed cases and as high as 23-44% in severe patients [25-29]. Studies have shown that the mortality was markedly higher in patients with cardiac injury than in patients without cardiac injury (51.2-59.6% vs 4.5-8.9%) [3, 30]. In addition, patients with underlying CVD are more likely to develop acute myocardial injury [26, 30]. Concurrent occurrence of underlying cardiovascular disease (CVD) and myocardial injury causes a significantly higher mortality than patients with underlying CVD but without cardiac injury, (69.4% vs 13.3%)[30], suggesting that myocardial injury played a greater role in the fatal outcome of COVID-19 than the presence of underlying CVD itself.

ACE2 is an important target for SARS-CoV and SARS-CoV-2 infection, and heart is one of the organs which express ACE2[31]. During the SARS epidemic, the SARS-CoV viral RNA was detected on autopsied human heart samples[32]. Similarly, Guido Tavazzi et al reported the first case of myocardial localization of SARS-CoV-2[33]. Furthermore, hiPSC-CMs are susceptible to SARS-CoV-2 Infection in vitro [8]. Those studies all suggested that SARS-CoV-2 may directly infect heart in human body through ACE2. However, the pathogenesis of SARS-CoV-2 infection-related acute myocardial injury is still unknown. Previous studies suggested that pathogen-mediated direct myocardial injury, acute systemic inflammatory response [1, 4] and low blood oxygen levels [6]may be the most critical causes of myocardial damage in COVID-19 patients. In this study, we described several specific mechanisms of cardiac injury caused by SARS-CoV-2 direct infection. We used R software and bioinformatics to deeply analyze the RNA-Seq dataset GSE150293, which compared the gene expression between hiPSC-CMs and SARS-CoV-2-infected hiPSC-CMs samples. The results identified 1554 DEGs, including 726 upregulated genes and 828 downregulated genes. 4-dimensional GO analysis showed that the upregulated DEGs were mainly involved in defense response, receptor complex, cytokine receptor binding, cytokine activity, chemokine activity, type I interferon signaling pathway, regulation of adaptive immune response, neutrophil chemotaxis, regulation of type 2 immune response, and CD4-positive α-β T cell cytokine production. Results suggested that SARS-CoV-2 received a strong defensive response from hiPSC-CMs. Similarly, GSEA, KEGG pathway and PPI function analysis shown that immune response correlative signal pathways were activated by SARS-CoV-2, including TNFα, IL6-JAK-STAT3, IL2-STAT5, NF-κB, IL17, Toll-like receptor, and lipopolysaccharide-mediated signaling pathway. 11 hub genes associated with inflammatory were identified based on PPI network, including IL6, CXCL8, TLR4, STAT1, IL1B, CXCL10, ICAM1, JUN, CCL5, CCL2 and CD44. Similar with previous studies [34, 35], SARS-CoV-2 infection activates multiple immune responses, including innate immunity (via Toll-like receptor signaling pathway), adaptive and type-2 immune response. Consistent with the inflammatory cytokines detected in blood of patients with COVID-19[1, 36], hiPSC-CMs produce and activate various of cytokines, including TNFα, chemokines, interleukins (IL), and interferons after SARS-CoV-2 infection. The massive cytokine release reflects an excessive immune defense, which may cause harmful heart damage. These findings complement our understanding of the human immune response mechanism triggered by SARS-CoV-2 [34]. These hub genes or pathways may be potential targets for preventing myocardial damage or even treating COVID-19. Similar to our study, Omar Pacha et al suggested IL17 is immunologically plausible as a target to prevent ARDS in COVID-19[37].

GSEA shown that hypoxia and apoptosis gene sets were positively enriched in CoV group, whereas oxidative phosphorylation, E2F targets, G2M checkpoint, myogenesis and MYC targets v1 gene sets were enriched with Mock group. Go analysis of BP shown muscle system process, muscle contraction, oxidative phosphorylation, striated muscle contraction and respiratory electron transport chain were significantly enriched with downregulated DEGs. Cell compositions, such as myofibril, contractile fiber, sarcomere, I band and inner mitochondrial membrane protein complex, were inhibited by SARS-CoV-2. In the MF group, SARS-CoV-2 inhibit actin binding, NADH dehydrogenase (ubiquinone & quinone) activity and structural constituent of muscle. KEGG pathway analysis shown thermogenesis, oxidative phosphorylation, cardiac muscle contraction, retrograde endocannabinoid signaling, and adrenergic signaling in cardiomyocytes were enriched with downregulated DEGs. Those results showed that SARS-CoV-2 infection caused respiration dysfunction, muscle contraction disorders, and cell cycle arrest in HIPSC-CMS. Excessive inflammatory response, respiration dysfunction and contraction disorders can promote each other and aggravated the myocardial injury. The analysis of PPI network shown that SRAS-CoV-2 inhibit the regenerative potency of hiPSC-CMs by inhibiting the metaphase/anaphase transition of mitotic cycle, suggesting that the down-regulated 4hub genes (CDK1/UBE2C/CDC20/AURKB) may be important genes for myogenesis. Further research of these hub genes may help promote cardiac function recovery in COVID-19 patients.

Furthermore, ISP analysis of downregulated DEGs shown megakaryocyte differentiation and thymus development were repressed by SRAS-CoV-2 infection. These mechanisms may be responsible for the significant reduction of platelet count [38] and peripheral blood T cells [36]in patients with severe COVID-19.

Similar to clinically reported cardiac complications in COVID-19 patients, CLINVAR human diseases analysis shown SARS-CoV-2 infection may lead to myocardial infarction[39, 40], cardiomyopathy[41] and limb-girdle muscular dystrophy. We suspect that some COVID-19 patients may have long-term cardiac insufficiency after recovery.

Overall, our study showed that SARS-COV-2 infection can induce a strong immune inflammatory response, reduce contractility, increase hypoxia, and induce apoptosis in cardiomyocytes. It suggested that SARS-COV-2 could directly infect cardiomyocytes in the body and cause viral myocarditis, resulting in myocardial injury. This conclusion explains the prevalence of myocardial damage in COVID-19 patients. It should be noted that the occurrence of myocardial injury may be a sign of SARS-COV-2 entering the circulation, which also explains the acute kidney injury is more common among patients with cardiac injury[3].

The current study has several limitations. The study included only one available data packet of 6 hiPSC-CMs samples in vitro, which could not fully reflect the effect of SRAS-COV-2 on the heart in vivo. Only bioinformatics analysis was conducted in this study, further clinical reports or in vitro studies are needed to support our conclusions.

## Author Contributions

XFL, LQL and LZ designed analyses and drafted the manuscript. XFL and LQL performed the bioinformatics analysis. LZ reviewed the manuscript and provided funding support. All authors contributed to the final manuscript.

### Funding

This study was funded by National Natural Science Foundation of China (Grant/Award Numbers: “81970723”)

### Conflict of Interest

The authors declare that they have no conflict of interest.

### Availability of data and Materials

Please contact corresponding author for data requests.

## Notes

### Competing Interest Statement

The authors have declared no competing interest.

